# CRISPR-Cas12a induced DNA double-strand breaks are repaired by locus-dependent and error-prone pathways in a fungal pathogen

**DOI:** 10.1101/2021.09.08.459484

**Authors:** Jun Huang, David Rowe, Wei Zhang, Tyler Suelter, Barbara Valent, David E. Cook

**Affiliations:** Department of Plant Pathology, Kansas State University, Manhattan, KS 66506-5502, USA

**Author notes:** Address correspondence to David E. Cook.

## Abstract

CRISPR-Cas mediated genome engineering has revolutionized functional genomics. However, basic questions remain regarding the mechanisms of DNA repair following Cas-mediated DNA cleavage. We developed CRISPR-Cas12a ribonucleoprotein genome editing in the fungal plant pathogen, *Magnaporthe oryzae*, and found frequent donor DNA integration despite the absence of long sequence homology. Interestingly, genotyping from hundreds of transformants showed that frequent non-canonical DNA repair outcomes predominated the recovered genome edited strains. Detailed analysis using sanger and nanopore long-read sequencing revealed five classes of DNA repair mutations, including single donor DNA insertions, concatemer donor DNA insertions, large DNA deletions, deletions plus donor DNA insertions, and infrequently we observed INDELs. Our results show that different error-prone DNA repair pathways resolved the Cas12a-mediated double-strand breaks (DSBs) based on the DNA sequence of edited strains. Furthermore, we found that the frequency of the different DNA repair outcomes varied across the genome, with some tested loci resulting in more frequent large-scale mutations. These results suggest that DNA repair pathways provide preferential repair across the genome that could create biased genome variation, which has significant implications for genome engineering and the genome evolution in natural populations.

## Introduction

The CRISPR-Cas9 (clustered regularly interspaced short palindromic repeats and CRISPR associated protein 9) genome editing platform has been widely used in multiple organisms including animals, plants, and fungi for functional genomics studies (1–5). The basic requirement for CRISPR-Cas genome engineering is a Cas endonuclease protein complexed with a single-guide RNA targeting a genomic region following a protospacer adjacent motif (PAM), such as NGG in the case of the commonly used SpCas9 protein from *Streptococcus pyogenes* (1,2). Another Cas effector, termed Cas12a (formerly named as Cpf1), is an alternative genome editing tool that has several unique features compared to Cas9 based effectors (6–12). For instance, Cas12a recognizes a T-rich PAM, which can be better suited for editing some genomic regions (7). Also, the RNase activity of Cas12a can process an array (single RNA molecule with multiple guide sequences) into multiple RNA molecules of single sequences, which allows more convenient multiplex genome engineering (6).

A critical component determining the outcome of genome engineering is DNA double-strand break (DSB) repair, mediated by endogenous DNA repair machinery (13). Proper repair of DNA DSBs, whether induced by Cas effectors or under natural conditions, is critical to maintain genomic stability, where repair failure can result in altered genome function and be potentially lethal (14–16). DNA DSB repair is mediated by two major pathways, canonical non-homologous end joining (C-NHEJ) and homology direct repair (HDR) (17–19). One of the major differences between C-NHEJ and HDR is the initial processing of DNA ends at a break site, where HDR requires extensive DNA end resection (i.e., enzymatic nucleotide removal from DSB site), which is inhibited by NHEJ (19,20). In the initial steps of C-NHEJ, the Ku70/Ku80 heterodimer interacts with broken DNA ends to inhibit resection (21), and recruits additional proteins to the site eventually repairing the DSB via DNA ligase IV (17,22,23). The C-NHEJ pathway does not rely on a homologous DNA template for repair, and commonly results in small insertions and deletions (INDELs) (17,24), but there are also examples of accurate NHEJ repair with and without DNA templates (25,26). For DNA DSB repair via the HDR pathway, template DNA with extended homologous sequences (typically >100 bp) are used for what is generally considered accurate repair (18). Two additional DNA DSB repair pathways, which also require end resection at DSB sites, are termed alternative end-jointing (a-EJ), and single strand annealing (SSA) (17,18,27). The a-EJ pathway is also referred to as microhomology-mediated end-joining (MMEJ), theta-mediated end-Joining (TMEJ), and has been called alternative NHEJ (A-NHEJ) depending on the system and report (28,29). While the three pathways involving end resection rely on homologous sequence for DSB repair, the length of homologous sequence used by a-EJ, SSA and HDR is different. The a-EJ repair involves annealing microhomologous sequences (typically 2-20 bp) and gap filling by DNA polymerase theta (Polθ) near the DSB (30), resulting in small insertions, deletions and templated insertions in mammalian and plant systems (17,31,32). The SSA pathway involves annealing with longer homologous sequences (>25 bp), often described to reside at longer distances from the DSB site and result in larger deletions as the result of removing 3’ non-homologous ssDNA via Rad1-Rad10 endonuclease (17,18,27,30).

Many questions remain for how the individual DNA repair pathways interact, such as their individual contributions to genome stability, their hierarchy for DSB repair, and variation in DSB repair pathways in microbial eukaryotes. There have been conflicting reports on the importance and role of a-EJ for repairing DSBs (17,33), however, clear evidence shows that a-EJ substantially contributes to DNA repair in zebrafish embryos (34), mouse cell lines (35), *Caenorhabditis elegans* (36), human cells lines (37) and the model plant *Arabidopsis thaliana* (38). Interestingly, while the identification of C-NHEJ independent DSB repair was first described in *Saccharomyces cerevisiae* (39), there are no reports of TMEJ in yeast or filamentous fungi. There are also clear differences for DSB repair across fungi, such as HDR being highly active in yeast *S. cerevisiae*, while C-NHEJ predominates DNA DSB repair in most filamentous fungi (40–42).

In this research, we developed efficient genome editing using Cas12a-based ribonucleoprotein (RNP) in *Magnaporthe oryzae* (synonym of *Pyricularia oryzae*), a filamentous fungal pathogen of monocots threating world food security (43–45). The use of CRISPR editing in fungi can increase the speed and efficiency of traditional gene replacement strategies, and the use of RNPs can alleviate problems related to cytotoxicity and off-target mutations (46–48). Surprisingly, we found that Cas12a editing in *M. oryzae* resulted in numerous mutants that contained severe DNA alterations at the targeted locus. Using long-read DNA sequencing and *de-novo* assembly, we confirmed at nucleotide resolution, multiple classes of DNA mutations, suggesting the involvement of different DNA repair mechanisms. The frequency of DNA repair outcomes after Cas12a editing were found to be locus-dependent across five tested loci. These results provide a first detailed report of variable DNA repair outcomes after Cas12a-RNP editing which have significant implications for natural and induced DNA repair in fungal genomes.

## Materials and Methods

### Fungal strains and incubation condition

*M. oryzae* field isolates O-137 (China) and Guy11 (French Guyana) were used as wild-types in this study (43,49). *buf1* mutants CP641 (gained from O-137 through spontaneous mutation) and CP281 (derived from weeping lovegrass pathogen 4091-5-8 through UV-mutagenesis) were used as a control in identifying the buff phenotype (Valent, unpublished data). JH7#1 and #2 are FK506 resistant mutants caused by a mutation in the *FKBP12* gene. The fungal cultures were maintained under light at 25 °C on OTA to observe mycelial color change. For high-molecular-weight DNA extraction and protoplast preparation, related mycelial plugs for different strains from OTA were cultured in liquid CM at 28°C, 120rpm for 3-4 days as described before (50).

### *In vitro* crRNA synthesis and LbCas12a RNP assembly

Oligos including T7 promoter (taatacgactcactatagg), LbCas12a direct repeat (taatttctactaagtgtagat), and 23-nt target sequences (Supplementary Table 2) were annealed and amplified to make the DNA template for *in vitro* RNA synthesis. HiScribe™ T7 High Yield RNA Synthesis Kit (New England BioLabs, catalog# E2040S) was used to make the crRNA/gRNA with the above prepared DNA template according to manufacturer’s protocol. Monarch® RNA Cleanup Kit (New England BioLabs, catalog# T2050L) was used to purify synthesized gRNA after DNase I (RNase-free) treatment (New England BioLabs, catalog# M0303S). 5 μg purified LbCas12a (New England BioLabs, catalog# M0653T) was incubated with equal molar purified gRNA (near 0.5 μg) at 25°C for 15 mins for RNP assembly.

### DNA donor preparation

pFGL821 (hygromycin selection, a gift from Dr. Naweed Naqvi; Addgene plasmid # 58223), and pFGL921 (G418 selection) (50) were used as DNA templates for amplifying DNA donor with related primer pairs (Supplementary Table 2). Long homologous sequences of SS flanked *HYG* DNA donor were amplified from O-137 genomic DNA and inserted into KpnI/XbaI and SalI/PstI sites in pFGL821 (Supplementary Table 2). Phusion® High-Fidelity DNA Polymerase (New England BioLabs, catalog# M0530L) was used for DNA donor amplification.

### Protoplast preparation and polyethylene glycol (PEG) mediated transformation

*M. oryzae* protoplast preparation was performed as described previously with minor modifications (51). The fungal mycelium was filtered and dried through 2-layer 61/2 in Disks Non Gauze Milk Filter papers, followed by addition of lysing solution (10 mg/mL Lysing Enzymes from *Trichoderma harzianum*, Sigma, catalog# L1412-10G, dissolved in 0.7M NaCl solution) and digestion at 30 °C with 70-80 rpm for 2-3 hours in the dark. After washing with 1xSTC (20% w/v Sucrose, 50mM Tris-Hcl pH=8.0, 50mM CaCl_2_ dissolved in water), the concentration of released protoplasts was adjusted to 8×10^6^-5×10^7^protoplasts/mL for transformation.

For protoplast transformation, RNP complexes (5 μg LbCas12a protein complexed with equal molar amount of crRNA/gRNA) and 3 μg DNA donor were mixed with 200 μl concentrated protoplast at room temperature for 20-25 mins. 1 mL 60% PEG solution (60% PEG4000, 20% w/v sucrose, 50 mM Tris-HCl pH=8.0, 50 mM CaCl_2_) was added to above mixture and incubated at room temperature for 20-25 mins. This was followed by incubation with 5 mL TB3 liquid medium at 28°C, 90rpm, for 10-18 hours. After overnight incubation, the fungal cultures were mixed with 50mL molten (near 50-60°C) TB3 solid medium containing 100 μg/mL hygromycin (Corning, catalog# 45000-806) or 300 μg/mL G418 (VWR, catalog# 97064-358). The fungal medium suspension was poured into a plate (150 x 15 mm), dried and then overlaid and cultured with another 50mL molten TB3 solid medium plus 200 μg/mL hygromycin, or 600 μg/mL G418 in dark condition at 28°C for 5-7 days. Potential fungal transformants were picked and sub-cultured on CM, OTA or RPA for further phenotyping and genotyping. CM was supplemented with 1μg/mL FK506 (LC laboratories, catalog# NC0876958), for testing the sensitivity to FK506.

### PCR genotyping

To test genotypes for the gene of interest, Q5® High-Fidelity DNA Polymerases (New England BioLabs, catalog# M0491) was used with 2 min extension time (technically up to 6 kb amplification) for the first-round genotyping with gene specific primer pairs (gene_F/R) (Supplementary Table 2). Upstream and downstream regions and *ACTIN* were amplified with *Taq* DNA Polymerase (New England BioLabs, catalog# M0273) (Supplementary Table 2).

### High-molecular-weight DNA extraction and library preparation for Nanopore sequencing

The DNA extractions for long-read nanopore sequencing were performed following the online protocol (https://www.protocols.io/view/high-quality-dna-from-fungi-for-long-read-sequenci-k6qczdw) (52). g-Tube (Covaris, catalog# 520079) was used for shearing high-molecular-weight DNA into 20kb fragments followed by purification with AMPure XP beads (Beckman, catalog# NC9959336). Nanopore sequencing library preparation followed the Native barcoding genomic DNA (with EXP-NBD104, EXPNBD114, and SQK-LSK109) protocol (https://community.nanoporetech.com/protocols/native-barcoding-genomic-dna/checklist_example.pdf). Eight barcoded DNAs from independent Cas12a edited strains were sequenced in two nanopore MinION platforms for 72 hours.

### Genome assembly and long-read mapping

Raw MinION fast5 files were transferred to fastq files by Guppy (version 3.4.4, https://nanoporetech.com/nanopore-sequencing-data-analysis) with the following parameters: --disable_pings --compress_fastq --flowcell FLO-MIN106 --kit SQK-LSK109. Adaptors were removed from basecalled reads by Porechop (version 0.2.4, https://github.com/rrwick/Porechop). Canu (version 1.9 and 2.2.1, https://github.com/marbl/canu) (53) was used for *de novo* genome assemblies with the following parameters: genomeSize=45m, minReadLength=1500. Based on the flanking sequences, the contigs with *BUF1* flanking sequences (including #7, #8 assemblies from Fig. 3d) were merged manually to scaffold based on the alignment result, when the contigs containing *BUF1* are not intact. The raw long-read mapping was performed with minimap2 (version 2.17-r941) (54). The mapping results were adjusted with samtools (55) with following parameter: samtools view -q 60 to reduce the duplicated mapping and visualized with the Integrative Genomics Viewer (IGV) (version 2.5.0) (56)

### Synteny analysis

The synteny plots between sequenced Δ*buf1* and wild-type (O-137) were generated with Easyfig (version 2.2.5) (57) by with the following parameter: Blast option_Min. length 30, Max. e Value 0.001. Each individual synteny plot was modified with Adobe illustrator.

### Statistical analysis

Fisher’s exact test was performed in RStudio (Version 1.2.5001) with function (fisher.test).

### Phylogenetic and domain analysis

The protein sequences of human Polq homologous were extracted from NCBI Genbank (https://www.ncbi.nlm.nih.gov/genbank/) and FungiDB (https://fungidb.org/fungidb/app/). MGG_15295 (*M. oryzae*), NCU07411 (*Neurospora crassa*), FGRAMPH1_01G23295 (*Fusarium graminearum*), NP_955452.3 (*Homo sapiens*), NP_084253.1 (*Mus musculus*), NP_524333.1 (*Drosophila melanogaster*), NP_498250.3 (*Caenorhabditis elegans*), AT4G32700.2 (*Arabidopsis thaliana*), XP_015619406 (*Oryza sativa*) and Pp3c5_12930V3.1 (*Physcomitrella patens*). The Neighbor-joining tree was made with MEGA X (58) with 1000 bootstrap value. TBtools (59) was used visualized the domain structure gained from pfam (http://pfam.xfam.org/).

## Results

### Cas12a ribonucleoprotein complex mediates efficient DNA editing

To test Cas12a RNP gene editing in *M. oryzae*, we designed two gRNA targeting the *BUF1* locus that codes for a trihydroxynaphthalene reductase required for fungal melanin biosynthesis (60). A DNA nuclease competent RNP comprised of purified LbCas12a protein (*Lachnospiraceae bacterium ND2006*) and *BUF1*-gRNA1 or -gRNA2 were transferred with donor DNA coding for the hygromycin resistance gene (*HYG*) into *M. oryzae* field isolate O-137 using protoplast transformation (Fig. 1a) (49). The donor DNA contained short (30 and 35 bp) flanking sequences at the ends, homologous to the *BUF1* locus, to direct microhomology-mediated end joining (MMEJ donor DNA integration following Cas12a DNA DSB) (Fig. 1b). Transformed protoplasts were recovered on hygromycin selection, where >50% displayed the mutant buff phenotype (i.e., orange/tan color) for both tested gRNAs. Control transformations with donor DNA alone exhibited wild-type hyphal pigmentation (Fig. 1c and Supplementary Fig. 1). To genotype and confirm Cas12a-mediated editing of *BUF1*, PCR was used to discriminate wild-type (~1.5 kb product) versus single-copy *HYG* donor insertion (~3.1 kb product) or larger visible PCR product corresponding to the integration of *HYG* DNA donor, which we refer to as a ‘simple insertion’ (Fig. 1b, d). The Cas12a-mediated DSB could also be repaired to create an INDEL resulting in a PCR product indistinguishable from wild-type, while hygromycin resistant transformants with wild-type pigmentation were presumed to have integrated the selectable marker at a secondary locus (Fig. 1d). Across experiments, the *BUF1* locus was edited at a rate of ~60% (14/23), but interestingly, all transformants that displayed the mutant buff phenotype generated with *BUF1*-gRNA1 (7/7), and almost half generated with the *BUF1*-gRNA2 (3/7), failed to produce a *BUF1* PCR product (Fig. 1e). The other 4/7 mutants generated with *BUF1*-gRNA2 produced a PCR band consistent with integration of a single copy of the *HYG* coding sequence (Fig. 1e). All recovered transformants displaying a wild-type hyphal color produced the anticipated wild-type sized PCR product (Fig. 1e). The *BUF1*-gRNA1 targeted an intron and could therefore have generated DNA mutants that failed to produce a visible phenotype. To assess this, transformants generated with *BUF1*-gRNA1 that had a wild-type hyphal color and wild-type PCR amplificon were sanger sequenced, which showed that all strains with normal pigmentation had wild-type *BUF1* sequence (Fig. 1e, f). We further confirmed these results by repeating the experiments in a different rice blast field isolate, termed Guy11 (43). These results using Guy11 were consistent with those observed in O-137, in which the majority of transformants (12/17) that had the mutant buff phenotype failed to produce a PCR product from the *BUF1* locus (Fig. 1g and Supplementary Fig. 2). We had anticipated that the majority of mutants would have a single donor integration mediated by the homologous sequence on the donor DNA, but this only occurred in ~29% (9/31) of the O-137 and Guy11 buff mutants. Furthermore, the majority (~71%, 22/31) of repair events resulted in the inability to generate a PCR amplicon, suggesting a severe DNA alteration following DNA repair. From these first experiments, out of a total of 41 hygromycin positive transformants generated using Cas12a RNP across the two *M. oryzae* strains, we did not recover any *BUF1* INDEL mutations as is commonly reported for CRISPR-Cas DNA editing (Fig. 1f, g).

**Figure 1.**
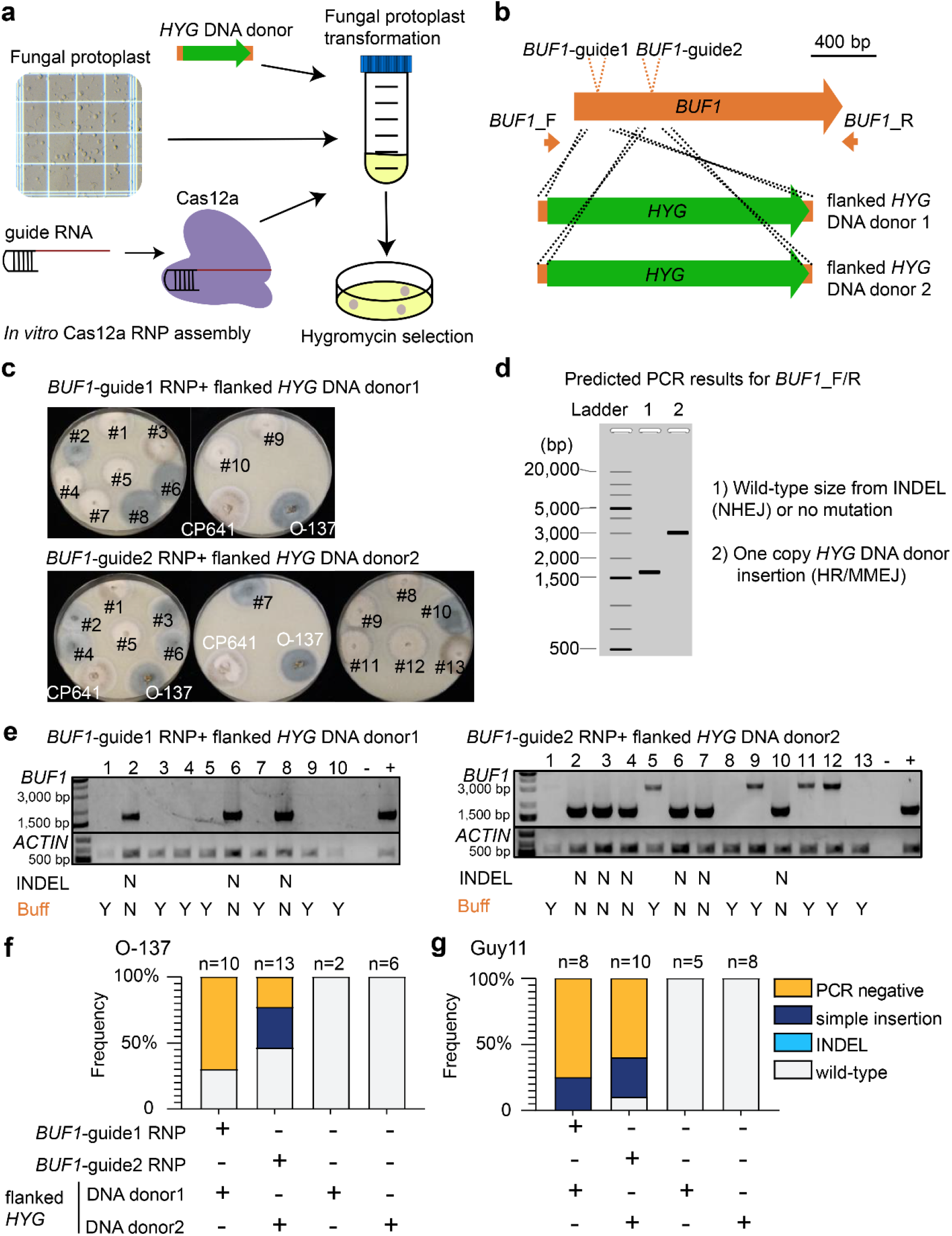
Cas12a RNP combined with short homology donor DNA mediates efficient gene editing in *BUF1* locus. (a) Schematic diagram of CRISPR-Cas12a RNP mediated genome editing through protoplast transformation in *M. oryzae*. (b) Illustration of two *BUF1* (MGG_02252, 70-15 MG8 annotation) gRNA design. Green rectangles indicate two different *HYG* DNA donor with flanked sequence homologous to the *BUF1* locus shown in orange. The location of PCR primer pair used for genotyping is shown (*BUF1*-F/R). (c) Representative phenotypes of hygromycin resistant transformants plated on OTA. The wild-type O-137 is shown (dark grey hyphae) and a previously characterized Δ*buf1* in O-137 (CP641) showing the buff phenotype. Individual transformed colonies showing wild-type and buff color hyphae are shown labeled with numbers (d) Diagram of expected results following PCR amplification from transformed lines. Ladder indicates the molecular weight ladder to determine product size, lane 1 (1) shows the expected size product for wild-type or small INDEL *BUF1* amplification, and lane 2 (2) shows the expected size product for *BUF1* amplification where a single copy of the *HYG* donor was inserted. (e) Representative genotyping results for the strains presented in the (c), the wild-type like PCR products from *BUF1* locus were purified and sanger sequenced to detect potential INDELs. INDEL N indicates there were no INDELs observed after sequencing. Buff Y indicates the strain displayed the buff mutant color, while Buff N indicates wild-type phenotype. Lane (−) indicates negative control (water) and (+) a positive control O-137 genomic DNA used for PCR amplification. A separate PCR to amplify a portion of *ACTIN* was used as a DNA extraction control. (f and g) The frequency summary of DNA DSB repair outcomes in O-137 and Guy11. The number of independent transformants (x) is listed (n=x).

### Extended homologous sequence is not required for donor DNA insertion at the Cas12a double-strand break site

Given that a majority of our mutants did not have a single donor DNA insertion, we speculated that the homologous sequence (30 and 35 bp) on the donor DNA was not required for Cas12a-mediated DSB repair. To test this, we performed the experiment again using Cas12a-RNP and the two gRNAs targeting *BUF1*, but used *HYG* donor DNA that lacked *M. oryzae* homologous sequence (no-homology *HYG*) (Fig. 2a). Transforming strain O-137, we obtained 61 hygromycin positive transformants, of which we could confirm 43 had a DNA mutation at *BUF1* (~70% editing efficiency). We found that ~60% (26/43) of the edited strains were buff colored and produced no PCR product, ~30% (13/43) were buff color and had a simple DNA insertion (roughly the size of a single copy of the donor DNA or other larger visible products) and ~9% (4/43) were wild-type color but had an intron INDEL from gRNA1 (Fig. 2b, c, f and Supplementary Fig. 3, 4). We additionally transformed strain O-137 with the two RNPs simultaneously (*BUF1*-gRNA1 and *BUF1*-gRNA2) and the no-homology *HYG* DNA donor (Fig. 2d), which resulted in a 75% editing frequency (12/16), where 42% (5/12) produced no PCR product, and 58% (7/12) had a simple insertion (Fig. 2d, e, f). Both gRNAs were used to repeat the experiments in the Guy11 strain producing a ~69% editing efficiency (20/29), where 45% (9/20) produced no PCR product, 55% (11/20) had a simple insertion, and none of the 29 hygromycin positive transformants had an INDEL at *BUF1* (Fig. 2g and Supplementary Fig. 5). For the dual RNP transformation ~42% (5/12) had a DNA mutation, where 20% (1/5) showed the PCR negative genotype and 80% (4/5) had a simple insertion (Fig. 2g and Supplementary Fig. 5). Through sanger sequencing randomly selected ‘simple insertion’ transformants, we found frequent ~2bp microhomology between no-homology *HYG* DNA donor and *BUF1* locus at the integration junction. (Fig. 2h). Additionally, to rule out that our observations are dependent on the *HYG* donor DNA and hygromycin selection, we performed the same experiments with a different donor DNA sequence coding for resistance to the drug G418 (Geneticin) (50). We again recovered transformants with the buff mutant phenotype and found that 70% (14/20) were PCR negative and 25% (5/20) had PCR amplification that suggested a simple insertion. We also recovered a single transformant (1/20) that carried a *BUF1* INDEL from guide2 through templated insertion (Supplementary Fig. 6).

**Figure 2.**
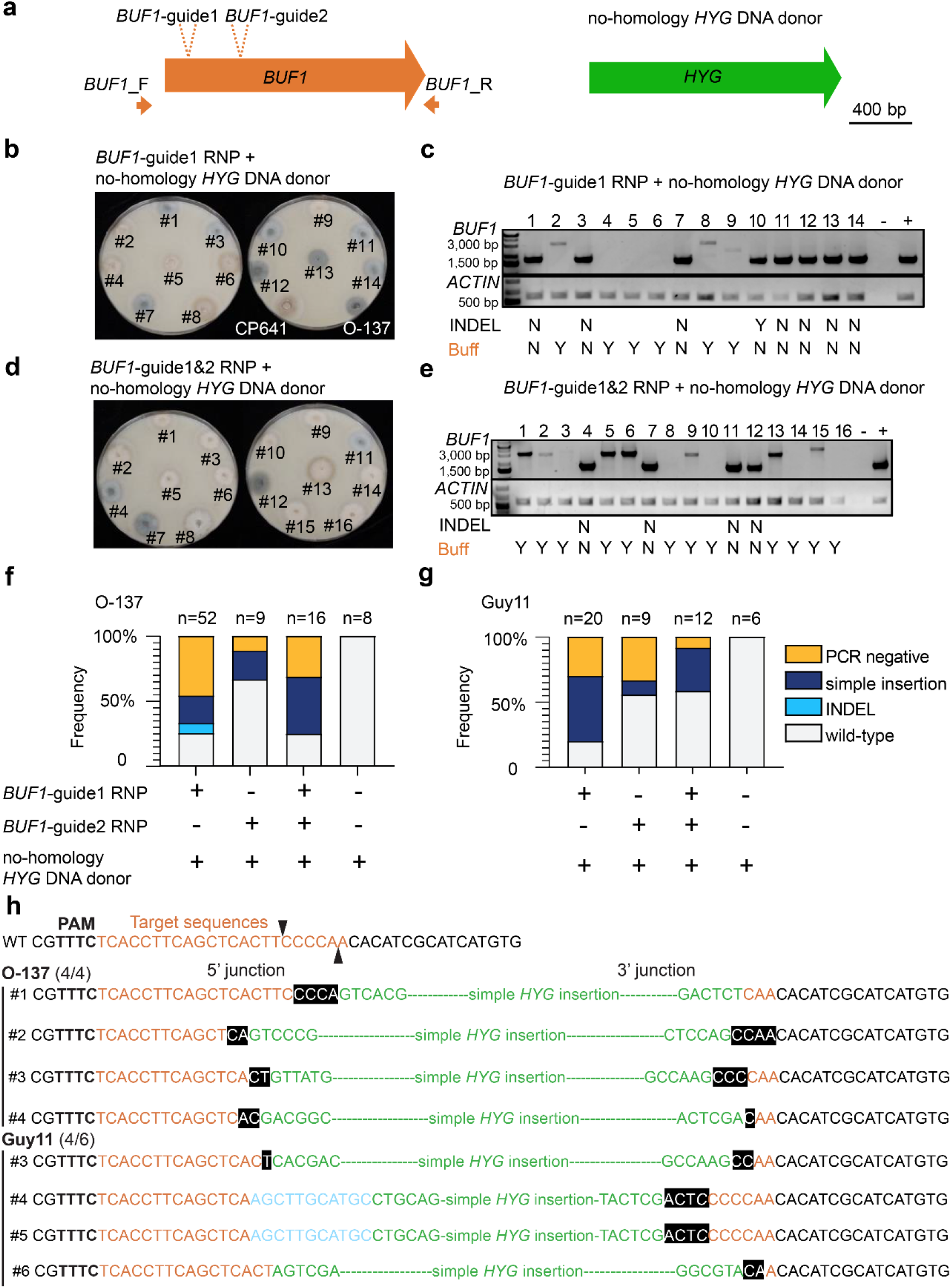
Donor DNA without homology sequences integrates at Cas12a target site. (a) No-homology *HYG* DNA donor with *BUF1*-guide1 or/and guide2 RNP were used for protoplast transformation. (b and d) Representative phenotyping result of the hygromycin resistant transformants from single or dual *BUF1* RNP targeting assays. The strains were plated on OTA for phenotyping. CP641 is a positive control (Δ*buf1*) for buff color hyphae, O-137 is the wild-type isolate used in the experiment. Individual transformed colonies showing wild-type and buff color hyphae are shown labeled with numbers. (c and e) Representative genotyping results for the strains presented in (b and d), the wild-type like PCR products from *BUF1* locus were purified and sanger sequenced to detect potential INDELs. Image labels are the same as described for figure 1. (f) The frequency of DSB repair outcomes in O-137. The number of independent transformants (x) is listed (n=x). The assays for *BUF1*-guide1 were counted from three independent rounds of transformation (rep1 to rep3). (g) The frequency of DSB repair outcomes in Guy11. (h) DNA sequence at the Integration junction of simple insertion mutants. The ratio to the right of the strain name is the number of mutants with at least 1 bp microhomology at the integration junction from the randomly selected simple insertion mutants sanger sequenced. Bold letters indicate PAM sequences; orange letters indicate target sequences; black triangles highlight the potential Cas12a cut site. Green sequences are from *HYG* DNA donor; White letters in black boxes highlight microhomology (i.e., shared sequence) between the *BUF1* locus and donor DNA; blue letters are sequence insertions of unknown source. Italicized letters indicate SNP.

**Figure 3.**
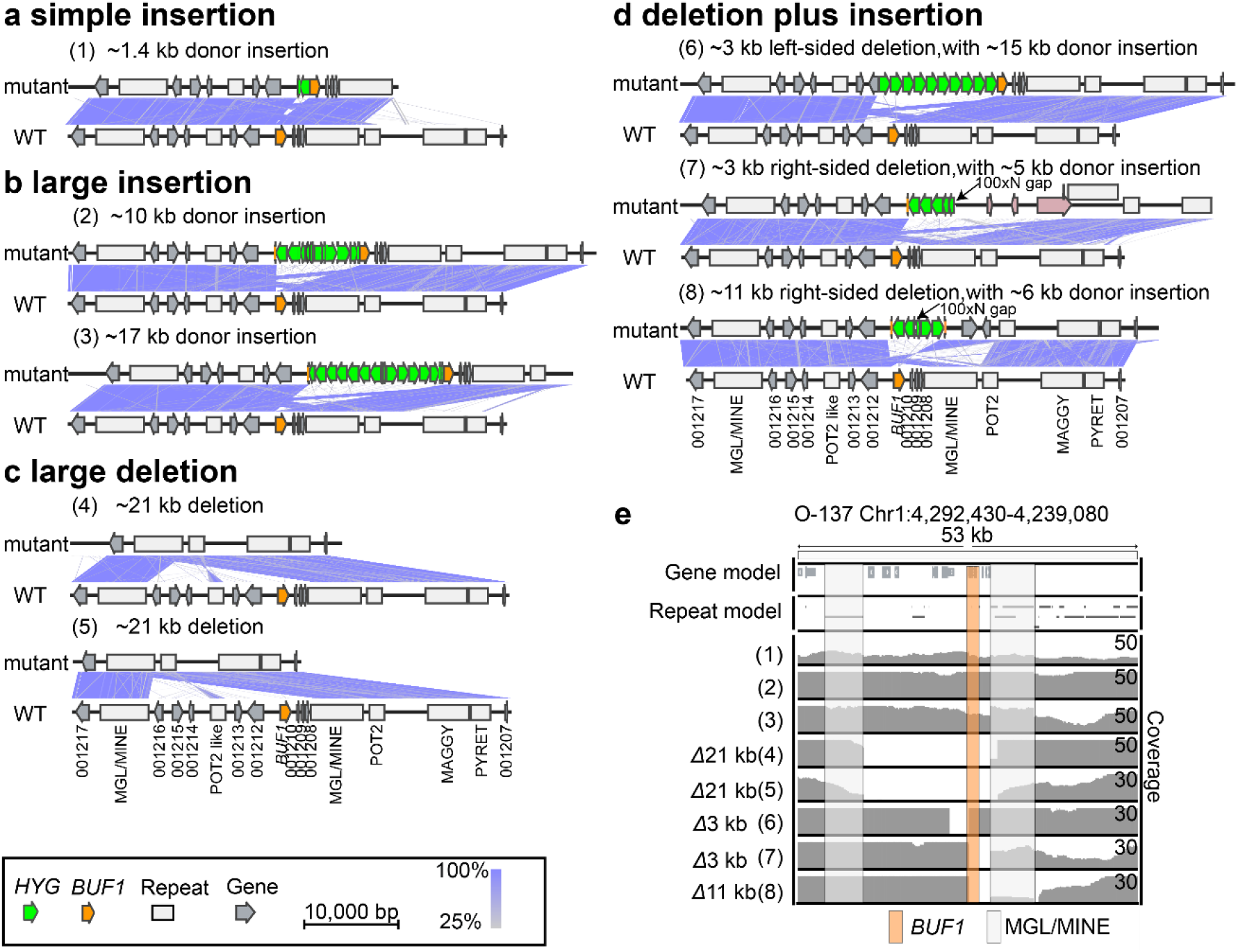
Long read assemblies reveal three non-canonical error-prone DNA repair outcomes. Microsynteny between assembled edited strains (top) and the wild-type (bottom) indicated by purple connecting bands ranging from 25 to 100% similarity as indicated by the color scale. Individual assemblies were classified into one of four mutation classes (a-d). (a) simple donor insertion, (b) large donor DNA insertion, (c) large DNA deletion, (d) DNA deletion plus donor insertion, where two assemblies were incomplete and required the identification of another contig to complete the locus. The merged region is indicated with an arrow and label 100xN gap. The pink labelled genes in (7) indicated three genes inserted from a *trans* locus. (e) Coverage of nanopore long-read mapping to the *BUF1* locus. Reads were mapped to the O-137 wild-type genome, where reads were filtered to remove low quality mapping (MAPQ < 60). The *BUF1* gene is highlighted with as a vertical orange box while the flanking repetitive DNA (MGL/MINE) are highlighted with a light grey box. Symbols are shown in the key to indicate the *BUF1* coding sequence (orange arrow), *HYG* coding sequence (green arrow), other coding sequences (grey arrow), and annotated repetitive DNA (grey box). Mutants labeled (1), (2), (3), (4), (5), (6), (7) and (8) indicate Rep1-Δ*buf1#*2, Rep1-Δ*buf1#*5, Rep1-Δ*buf1#*6, Rep1-Δ*buf1#*4, Rep4-Δ*buf1#*5, Rep4-Δ*buf1#*1, Rep1-Δ*buf1#*10 and Rep4-Δ*buf1#*13 mutants respectively (Supplementary Table 1).

From these experiments using a donor DNA with no-homologous sequence, we conclude that i) donor DNA does not require extended homologous sequence to resolve Cas12a-mediated DSB; ii) non-homologous donor DNA can integrate at DSB sites at a reasonably high frequency (40% across all experiments, 40/100) ; iii) more severe DNA alterations resulting in no PCR products are common (55% across experiments, 55/100); iv) INDELS were not common from these experimental conditions (5% across experiments, 5/100); v) targeted DNA mutation at *BUF1* is dependent on the Cas12a RNP complex, as 46 transformants obtained using no-homology *HYG* or *G418* donor DNA alone (i.e., in the absence of Cas12a RNP) did not cause the buff phenotype and among 26/46 that were PCR tested, none showed distinguishable *BUF1* DNA mutations (Fig. 2f, g and Supplementary Fig. 3d, 5d and 6c).

### Long-read sequencing and *de novo* assembly resolve genotypes following Cas12a-mediated DSB repair

We sought to further understand what DNA mutation occurred in the roughly 50% of buff mutants that failed to produce a PCR product at the *BUF1* locus. Eight O-137 derived buff mutants from the Cas12a RNP and no-homology *HYG* DNA donor transformation were selected for high-molecular-weight DNA extraction, nanopore sequencing and *de novo* assembly (Δ*buf1#2*, −*#4*, −*#5*, −*#6* from rep 1 in Fig. 2c, Δ*buf1#10* from Fig. 2e, Δ*buf1#1*, −*#5*, −*#13* from rep 4 in Supplementary Fig. 7a). All eight sequenced strains displayed the mutant buff color, where seven produced no PCR product, and one transformant (Δ*buf1#2* Fig. 2c) produced the ~3.1 kb PCR product we inferred to be a simple insertions of single-copy donor DNA. The eight strains were sequenced to an average depth of 52x and yielded highly contagious assemblies (average N50 of 3.29 Mb) (Supplementary Table1). These assemblies allowed for base pair resolution interrogation of *BUF1* DNA alterations. Consistent with PCR genotyping, the transformant thought to have a simple donor DNA insertion (Δ*buf1#2* Fig. 2c) indeed had an almost full copy of the hygromycin coding sequence (1,328 bp) plus an additional hygromycin fragment (140 bp) at the Cas12a cut site (16 bp after the PAM sequence) based on the long-read assembly (Fig. 3a, mutant 1). The insertion was nearly scar free, with the junction sequence showing 2 and 3 bp of microhomology at the 5’ and 3’ ends respectively, and only two bases pairs were deleted at the Cas12a endonuclease site (Supplementary Fig. 8). The results from the other seven assemblies were grouped into one of three categories, namely, large insertion, large deletion, and deletion plus insertion (Fig. 3b, c, d). Two mutants (Δ*buf1#5*, −*#6* from rep 1) each contained large insertions of concatemer *HYG* donor sequence, including promoter, coding, and terminator sequences, totaling 10 and 17 kb insertions respectively (Fig. 3b, mutant 2, 3). Not all the *HYG* DNA fragments were intact, and the coding sequences were in both the forward and reverse orientation (Fig. 3b). The insertion junction for both large insertion mutations had 2 bp of microhomology at the 5’ end, and no homology at the 3’ end, where in one mutant there was an error-free insertion at the locus, while the other mutant had a 17 bp deletion at the 3’ end (Supplementary Fig. 8).

Assemblies from two *BUF1* mutants (Rep1-Δ*buf1#4* and Rep4-Δ*buf1#5*) identified that same ~21 kb deletion around the *BUF1* locus, where *BUF1* and eight additional genes were all deleted (Fig. 3c, mutant 4, 5). Interestingly, the assemblies indicated the deletions took place between similar flanking non-LTR retrotransposons, which appear to be nested insertions of a LINE element, termed MGL, inserted into a hybrid LINE element termed MINE (61,62) (Supplementary Fig. 9). The deletions suggest that homology between the two elements was used to resolve the break, potentially through the SSA pathway, resulting in a single retrotransposon copy and the 21 kb deletion (Fig. 3c, mutant 4, 5, Supplementary Fig. 9). The *HYG* coding sequence for these two mutants was identified at independent loci on other chromosomes (Supplementary Fig. 10 a, b). In mutant 4, the HYG insertion was a large concatemer of ~20 kb, while the other deletion mutant had two *HYG* copies inserted (Supplementary Fig. 10 a, b). To confirm the assembly-based deletion, a ~6.7 kb PCR product that spanned the break resolution junction (MGL/MINE) was amplified in mutant 4, which failed to amplify a product in the wild-type as expected (Supplementary Fig. 11).

The remaining three mutants (Rep4-Δ*buf1#1*, Rep1-Δ*buf1#10* and Rep4-Δ*buf1#* 13) had both *BUF1* locus deletions, ranging in size from 3 to 11 kb on either the 5’ or 3’ side of the Cas12a endonuclease site, along with large insertions of concatemer donor DNA (Fig. 3d, mutant 6,7,8). The assemblies for mutants 7 and 8 did not completely resolved the *BUF1* locus in a single assembled contig. Here, we identified two contigs in both mutant assemblies with sequence homology to the *BUF1* locus, which were joined and analyzed. Interestingly, these results indicated the insertion of additional genomic DNA from other regions of the genome, including the insertion of coding sequences (Fig. 3d). As noted for the other mutation types, we also observed microhomology (2-3 bp) at three of the four integration junction sites (Supplementary Fig. 12). To support the assembly results, nanopore long-reads were mapped to the O-137 reference. Here we find a correspondence between the assembly-based deletions and a loss of read-coverage to the O-137 genome (Fig. 3e). These results support that large depletion and deletion plus insertion mutants lost DNA corresponding to the *BUF1* locus consistent with the *de novo* assemblies. Given that roughly half of the identified buff mutants across our experiments failed to produce PCR products, we were interested to use the assembly identified genotypes (large insertion, large deletion, deletion plus insertion) to screen the previously PCR negative transformants. For this, we designed primer pairs to amplify small fragments at the 5’ upstream and 3’ downstream of the *BUF1* locus (Supplementary Fig. 13a). Our results showed that 74% (23/31) of O-137 derived PCR negative transformants with no-homology *HYG* DNA donor and *BUF1* RNP had a large insertion (i.e., both 5’ and 3’ amplified PCR products), 9.6% (3/31) had a large deletion (i.e., both 5’ and 3’ PCR failed) and 16% (5/31) had a deletion plus insertion (i.e., PCR amplified product at either 5’ or 3’ end) (Supplementary Fig. 3a, b, 11b, d). We confirmed these results by also genotyping the transformants generated using the Guy11 derived strain, where we found that 80% (8/10) had a large insertion, 10% (1/10) had a large deletion and 10% (1/10) had a deletion plus insertion (Supplementary Fig. 13c, d).

### DNA repair outcomes following Cas12 editing differ between multiple loci

The *BUF1* locus has been characterized as unstable (60,63), and we were interested to understand if the unexpected DSB repair outcomes found at *BUF1* were representative of other loci in the genome. Wild-type *M. oryzae* is sensitive to the drug FK506, while disruption of the corresponding receptor, *FKBP12*, causes insensitivity to FK506 (Supplementary Fig. 14) (9,64–66). Therefore, we targeted *FKBP12* using Cas12a RNP and two separate gRNAs (*FKBP12*-guide1 and-guide2) and utilized sensitivity to FK506 to identify mutants (Fig. 4a). In order to compare the results to those from *BUF1, FKBP12* editing included the no-homology *HYG* DNA donor and hygromycin selection (Fig. 4b). We obtained 62 hygromycin resistant colonies, of which 13 (~21% editing efficiency) were insensitive to FK506 and presumably carried a mutation at the *FKBP12* locus, while none of the 20 no-homology *HYG* DNA alone transformants showed FK506 resistance (Fig. 4b and Supplementary Fig. 15). The same PCR amplification strategy was used to genotype the FK506 insensitive mutants, and we found that ~38% (5/13) had a simple insertion, ~54% (7/13) had a large insertion, and ~8% (1/13) had a deletion plus insertion (Fig. 4a, c and Supplementary Fig. 15a, b). None of the FK506 insensitive strains had an obvious wild-type sized PCR product (i.e., no INDELs) and the PCR genotyping indicated that no large deletions took place (Fig. 4a, c and Supplementary Fig. 15a, b).

**Figure 4.**
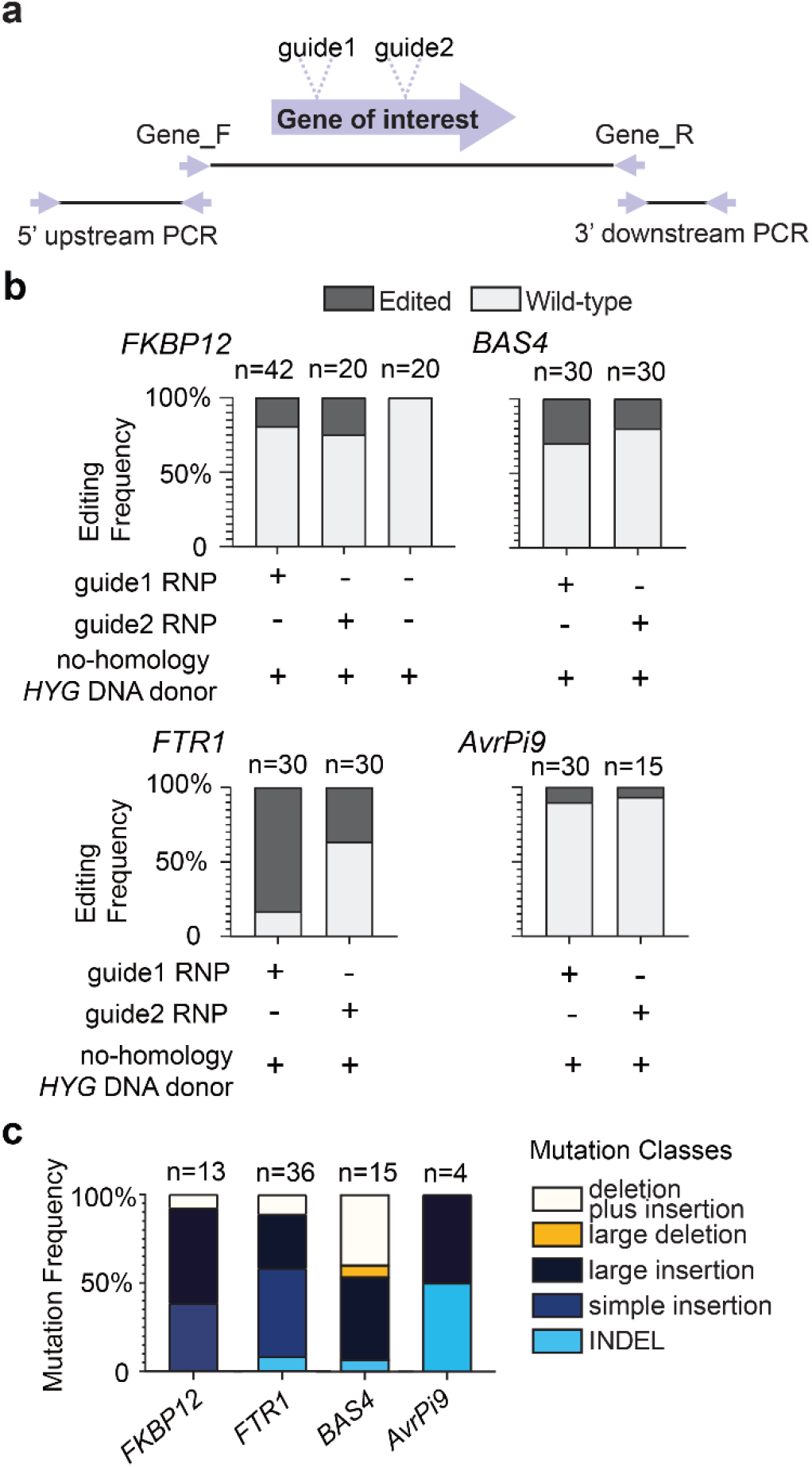
DNA repair outcomes following Cas12a editing differ among multiple loci. (a) Schematic illustration for CRISPR-Cas12a editing experiments. Two different guides were tested for each locus. Gene_F/R primer pairs were used for a first round of genotyping to determine the type of mutation present in a transformant. PCR negative strains were subsequently genotyped using the 5’ upstream and 3’ downstream primer pairs per gene to further genotype the occurrence of large-scale DNA alternations. (b) The editing frequencies at *FKBP12* (MGG_06035), *FTR1* (MGG_02158), *BAS4* (MGG_10914), and *AVRPi9* (MGG_12655). The MGG numbers based on the 70-15 assembly are provided for clarity (c) The mutation frequencies at *FKBP12*, *FTR1*, *BAS4* and *AVRPI9* are shown and indicated by the color key to the right. For (b) the number of independent transformants genotyped (x) is listed (n=x), while (c) lists the number of independent mutants from (b). Transformations using guide1 RNP, and guide2 for *FTR1* and *BAS4*, were performed twice independently, while guide2 for *FKBP12 and AvrPi9* were tested once.

We additionally tested three other loci, one coding for an annotated plasma membrane iron permease (*FTR1*) , which is 50 kb away from the *BUF1* locus, an apoplastic secreted protein *BAS4*, and an avirulence protein *AVRPi9*, both of which are presumed to help *M. oryzae* facilitate host infection (67–69). Two independent gRNAs were designed for each of the three loci, which were transformed as Cas12a-RNPs with the no-homology *HYG* DNA donor (Fig. 4b). From editing *FTR1*, 60 hygromycin resistant transformants were selected and genotyped. This gene had a high editing efficiency at 60% (36/60), and of the strains carrying a mutation, we found that ~8% (3/36) had an INDEL, 50% (18/36) had a simple insertion, ~31% (11/36) had a large insertion, ~11% (4/36) had a deletion plus insertion and no large deletions were recovered (Fig. 4b, c and Supplementary Fig. 16). The editing efficiency at *BAS4* was 25% (15/60), where ~6% (1/15) of the mutants had an INDEL, ~46% (7/15) had a large insertion, ~6% (1/15) resulted from a large deletion, 40% (6/15) from a deletion plus insertion (Fig. 4b, c and Supplementary Fig. 17). Interestingly, no simple insertions were recovered from the *BAS4* transformants using either of the two guides, despite simple insertion mutants being commonly found for the *BUF1* (30%), *FKBP12* (38%), and *FTR1* (50%) loci. Also of note, *BAS4* editing resulted in near half of the identified mutants having a deletion plus insertion or large deletion, which was not observed for other tested loci. The editing efficiency at the *AVRPi9* locus was only ~9% (4/45), and of these four mutants, two resulted from a large insertion and the other two lines carried INDELs (Fig. 4b, c and Supplementary Fig. 18). While we did not exhaustively test gRNA at all loci, the results using two independent gRNA at each locus suggests that editing efficiency is not the same across the genome, which has been reported in other organisms (70). An unexpected and novel finding, however, was that the spectrum of DNA mutations resulting from DSB repair did not occur at equal proportions for the tested loci. Indeed, formal testing of the different edited loci into the five classes of DNA mutations indicated that the highly significant association between the loci and DNA mutation outcomes, indicating a non-random association between repair mutation outcome and specific loci (Fisher’s exact test, p-value=0.006121).

### Locus-dependent DNA mutation frequency still occurs under a different editing scheme

Our initial editing experiments strongly favored donor DNA integration at the Cas12a-edited locus because of the induced DSB and selection on hygromycin. Given that our method was used across all edited loci, any bias should be similar among the loci. However, to further assess our results, we devised a second editing scheme. Here, we developed an assay where the donor DNA for selection was targeted to a separate non-coding locus, termed second-site (SS), by transforming with two distinct RNPs at once (Fig. 5a). This approach should capture DSB repair outcomes for the locus of interest (i.e., primary-site), without the need for donor DNA integration at the target locus, thereby decoupling donor DNA integration and selection from DSB repair outcomes. For these experiments, the *HYG* donor DNA contained long (730 bp and 518 bp) homologous sequence to the second-site targeted by Cas12a, and a separate RNP targeting the primary-site (Fig. 5a). Using this approach, we found high editing efficiency for the *BUF1* locus (89%, 25/28) (Fig. 5b), which was similar to the earlier results obtained (~77% with *BUF1*-guide1). The proportion of the five types of DNA mutations at *BUF1* were 28% (7/25) INDELs, 16% (4/25) simple insertion, 24% (6/25) large insertion, 32% (8/25) large deletions, while no deletion plus insertions were detected (Fig. 5c and Supplementary Fig. 19). These editing experiments returned substantially more INDEL mutations at *BUF1* than initially observed, consistent with the donor DNA integrating at the SS locus and not requiring it for DSB repair. Interestingly, the majority of recovered mutants still indicated either donor DNA insertion or large DNA deletion at the *BUF1* locus. The editing efficiency for the *FKBP12* locus was low, with only one mutant recovered from 41 transformants (Fig. 5b). The editing efficiency for this locus was also low for the single RNP assay, but it was not clear why it was even lower for this SS editing scheme. The one *FKBP12* mutant contained a large insertion at the locus (Fig. 5c and Supplementary Fig. 20). For the *BAS4* locus, we obtained ~33% editing efficiency (8/24), of which, 25% (2/8) were INDELs, 12.5% (1/8) were large insertion, 37.5% (3/8) were large deletions, and 25% (2/8) were deletion plus insertions, (Fig. 5b, c and Supplementary Fig. 21). Similar to initial single RNP assay, more than half of the *bas4* mutants were large deletion or deletion plus insertion mutations. The *FTR1* locus had a lower editing efficiency for the second-site assay compared to the initial single RNP editing with gRNA1 (38% versus 83%, respectively). Despite the reduction in editing efficiency and increased INDELs, the proportion of DNA mutation outcomes was quite similar between the two editing schemes, where the second-site assay showed 33% (4/9) INDELs, 33% (2/9) simple insertions, 22% (2/9) large insertions, and ~11% (1/9) deletion plus insertion mutations (Fig. 5b, c and Supplementary Fig. 22). In neither experiment did we recover large deletion mutants at *FTR1*. We additionally genotyped the second-site for each mutant strain and found PCR negative results for the second-site in most of cases (Supplementary Fig. 19, 20, 21 and 22). This suggested that the addition of long homologous sequences did direct the donor DNA to the second-site, but it did not provide precise homology directed repair in the form of a single donor DNA insertion. As controls, the second-site was also edited with Cas12a SS-RNP and second-site flanked *HYG* DNA donor in the absence of the primary site RNP and similar PCR negative results were found (Supplementary Fig. 23a). Also, flanked *HYG* donor DNA was transferred alone, without RNPs, and hygromycin resistant transformants did not show the buff mutant color or have FK506 resistance, nor did the transformants produce aberrant PCR products when amplifying the second-site locus, showing the mutations at these two primary sites, and the second-site were dependent on Cas12a-mediated DSB induction (Supplementary Fig. 23b). These results show that under a different editing scheme, which did not require donor DNA integration at the editing site of interest, we still observed a biased DNA mutation frequency following DSB repair at the tested loci.

**Figure 5.**
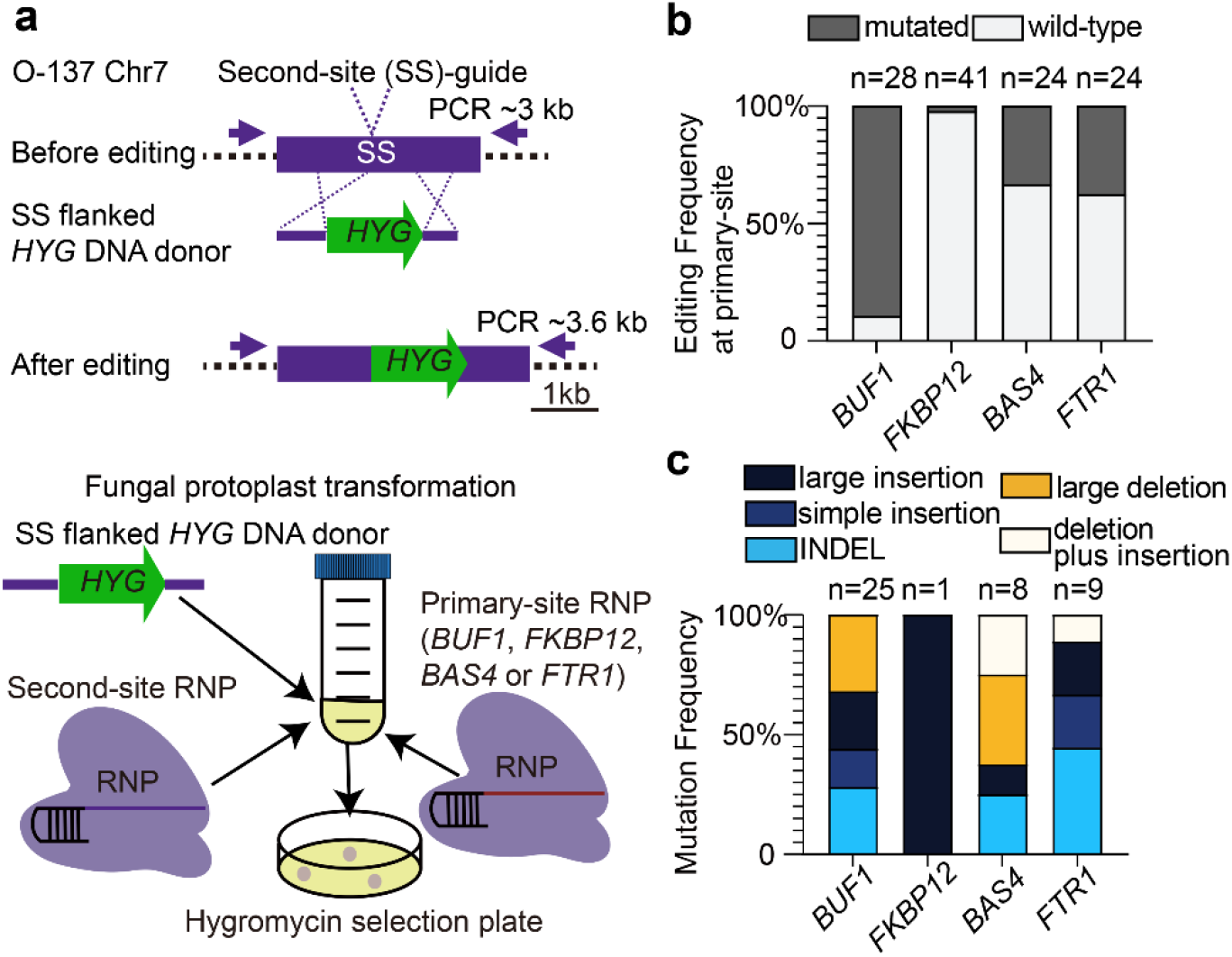
Distinct DSB repair pattens following donor insertion at a second integration site. (a) Schematic illustration of Second-site (SS) insertion assay. The *HYG* coding sequence was directed to a second site through long homologous sequence located in a non-coding region on chromosome 7 (Chr7:1,520,862-1,523,872). Fungal protoplasts were simultaneously transformed with two RNPs, one targeting the second-site and the other targeting the locus of interest. (b) The editing frequencies for the four tested loci following *HYG* selection. The proportion of mutated to wild-type transformants is indicated by dark and light grey shading respectively. (c) The frequency of distinct DNA repair outcomes for the four tested loci, colored according to the respective DNA mutation class. The number of independent transformants (b) and independent mutants (c) is listed (n=x). All the transformations were performed twice independently.

## Discussion

CRISPR-based genome engineering has accelerated functional genomic studies by providing a flexible platform to rapidly modify DNA and probe basic cell and molecular biology (1,7). Here we describe the development of an efficient and robust approach to modify specific loci in fungi using the Cas12a nuclease, delivered as an RNP. The use of RNPs for genome editing has the advantage of not requiring the integration and continual expression of the CRISPR-Cas system (48,71,72). This can overcome cytotoxicity, a reported attribute of continual expression of Cas nucleases in some systems including fungi, and is especially helpful for asexually reproducing fungi, where the CRISPR-Cas DNA cannot easily be removed through crossing (46,73,74). The strains generated in this study were made with a high success-rate, contain heritable mutations, and lack coding sequence of the CRISPR-Cas platform. We anticipate the described approach to deliver Cas12a RNPs to fungal protoplasts will be fungal species-agnostic and provides a rapid approach to generate gene disruption mutations, especially for recalcitrant loci (e.g., pathogen effectors) (75,76).

Our results highlight an unresolved question at the interface of genome evolution and genome biology, which is how does hierarchy and cross-talk between endogenous DNA DSB repair pathways influence genome variation? Using a combination of PCR, sanger sequencing and long-read sequencing-based assemblies, we show at base pair resolution how Cas12a-induced DNA DSBs can be variably repaired to generate a spectrum of DNA mutations. We observed INDELs, simple donor DNA insertions, large concatemer DNA insertions, large genomic deletions, and deletion plus insertion events, some of which resulted in drastic mutations at the targeted loci. It is difficult to determine the exact DNA repair pathway that resulted in each specific DNA mutation in the present work, however, there are mutation signatures of specific pathways that collectively suggest our mutation results were caused by at least three separate DNA repair pathways. This is based on the following i) we observed small insertion/deletion (INDEL) mutations with random sequence (i.e., not *cis*-template), which is a well characterized signature of C-NHEJ DSB repair dependent on Ku70/80 and Lig4 (18) ; ii) we observed a few INDELs that contain *cis-*templated insertions, which have been described as a hallmark for MMEJ (31,37); iii) we frequently observed microhomology between the genome and inserted donor DNA, highlighting the importance of sequence homology in DSB repair in our experiments. Microhomology was also found in the breakpoints of fused chromosomes in the human fungal pathogen *Cryptococcus deuterogattii* (77). The majority of junction sequences from our editing experiments did not contain non-templated insertions as has been reported for NHEJ knock-in insertions (25), and the insertions did not share the same genomic ends, a sign of DNA end resection. These DNA mutation patterns are signatures of MMEJ, and similar to those reported using staggered Cas9 editing in mouse (78), but the possibility that NHEJ is also involved cannot be ruled out. It should be noted, the mechanism of so-called microhomology mediated end-joining (MMEJ) is not well characterized in filamentous fungi. The a-EJ pathway described to be dependent on DNA polymerase theta (Pol θ) (i.e., TMEJ) has not been described in fungi, and *M. oryzae* lacks a clear homolog containing the DNA polymerase domain (Supplementary Fig. 24); iv) we identified large genomic deletions that resolved between two repetitive elements that reside 16 kb and 2 kb away from the induced DSB site, requiring extensive DNA end resection and the use of long homologous sequence for resolution, hallmarks of SSA (15,27). Similar large deletions between repetitive sequences have been reported in the protozoan parasite *Leishmania* (79). The large deletion mutants we observed suggest the potential importance of repetitive elements and SSA in fungal genome repair and evolution, however, the SSA pathway in filamentous pathogens is also not well documented. Based on these DNA mutation signatures, we can conclude that multiple DNA repair pathways were used to resolve the Cas12a-induced DSBs.

Along with demonstrating that multiple DNA DSB repair pathways were used to resolve the Cas12a-induced DSBs, we observed that these pathways were used at unequal frequencies across the tested loci (i.e., locus-dependent frequencies). This suggests a locus-specific hierarchy for DNA repair pathways, which to our knowledge has not been previously reported in fungi. It is unclear what factors may contribute to locus-dependent DNA repair preferences. Physical and genomic features such as chromatin structure, chromosome location, repetitive elements and the cell cycle have been reported to affect the outcome of DNA repair in other model systems (13,17,80–82). Demonstrating locus-dependent DNA repair is important because such a hierarchy would influence the evolutionary trajectory of different genomic regions. In natural systems, preferential repair of DSBs by different DNA repair pathways could create biased DNA variation and create genomic regions with accelerated evolution. There are numerous reports of compartmentalized genome evolution in filamentous pathogens, often referred to as two-speed genome evolution (83), and variation for DNA DSB repair could be a major driver of this phenomena. Indeed, detailed analysis of DNA translocations and inversions between strains of *Verticillium dahliae*, a soil-borne wilt causing pathogen, found that chromosome re-arrangements co-localize with homologous sequence, often transposable element DNA (84). Repetitive DNA can serve as templates for homology-based repair, such as MMEJ or SSA. The occurrence of variable DNA repair reported here may influence adaptive genomic regions in *V. dahliae* that are associated with a unique chromatin profile (81,85). In *M. oryzae*, we saw low editing efficiency at the *AvrPi9* locus, which has proven to be a stable R-Avr interaction for breeding blast resistant rice (67,86). It will be interesting to understand if DNA repair bias at the *AvrPi9* locus contributes to the durability of this interaction. Also in *M. oryzae*, the presence of mini-chromosomes that are sequence varaible between strains, and contain duplicated sequences with core chromosomes, may be impacted by DNA repair variation (44).

It is unlikely the observations reported here are specific to *M. oryzae* or our experimental setup. Genome editing in other filamentous fungi, such as *Sclerotinia sclerotiorum* and *Aspergillus fumigatus*, reported unexpected genotyping results in which the target loci were PCR negative (87–89). Further TAIL-PCR and Illumina sequencing suggested that vector sequences and many uncharacterized sequences were inserted in the target loci in the case of *S. sclerotiorum* (87). Those experiments used the Cas9 effector and different fungi, suggesting our results are not specific to Cas12a or *M. oryzae* (87–89). It should also be noted, large-scale on-target mutations created by CRISPR are easily missed without comprehensive genotyping or *de novo* assembly such as employed here, and are likely heavily under reported. Also, our repair outcomes are different than off-target editing often discussed for CRISPR research. Off-target editing refers to Cas nucleases binding and cutting DNA at unintended loci in the genome due to sequence homology (i.e., low-fidelity editing). Our results suggest that on-target DNA mutations following CRISPR DSB induction are varied and complex, consistent with recent reports in mammalian systems reporting extensive on-target mutations, including large deletion, complex rearrangements, and plasmid insertions (90–92). These results underscore the need to further understand how CRISPR-based genome engineering interacts with endogenous DNA repair mechanisms, and more importantly, how specific DNA repair mechanisms impact genome evolution in filamentous pathogens.

## Data availability

The *de novo* assemblies of Cas12a-edited strains from nanopore sequencing have been deposited in the WGS Genome of the National Center for Biotechnology Information (NCBI) under the BioProject accession No. PRJNA753862. The base called nanopore reads after adapter removal have also been deposited under the same BioProject accession No. PRJNA753862 with the Short Read Archive (SRA) accession No. SRR15459267 to SRR15459274.

## Acknowledgments

The authors would like to thank Dr. Qiang Wang (Auburn University), Dr. Sanzhen Liu and Dr. Huakun Zheng (Kansas State University) for helpful discussion during the preparation of the manuscript. J.H. would like to thank Haolang Jiang (Fujian Agriculture and Forestry University) for helpful suggestions for fungal transformation and Melinda Dalby, Nathan Ryan and Dr. Velazhahan Rethinasamy (Kansas State University) for technical support. The authors thank computational support from the Beocat High-Performance Computing cluster at Kansas State University.

